# Omicron-specific naive B cell maturation alleviates immune imprinting induced by SARS-CoV-2 inactivated vaccine

**DOI:** 10.1101/2024.05.13.594034

**Authors:** Ayijiang Yisimayi, Weiliang Song, Jing Wang, Fanchong Jian, Yuanling Yu, Xiaosu Chen, Yanli Xu, Ran An, Yao Wang, Jing Wang, Haiyan Sun, Peng Wang, Lingling Yu, Fei Shao, Ronghua Jin, Zhongyang Shen, Youchun Wang, Yunlong Cao

**Affiliations:** Biomedical Pioneering Innovation Center (BIOPIC), School of Life Sciences, College of Chemistry and Molecular Engineering, Peking University, Beijing, P.R. China; Changping Laboratory, Beijing, P.R. China; Institute for Immunology, College of Life Sciences, Nankai University, Tianjin, P. R. China; Beijing Ditan Hospital, Capital Medical University, Beijing, P.R. China; Organ Transplant Center, NHC Key Laboratory for Critical Care Medicine, Tianjin First Central Hospital, Nankai University, Tianjin, P. R. China; Institute of Medical Biotechnology, Chinese Academy of Medical Science & Peking Union Medical College, Kunming, P. R. China; Peking–Tsinghua Center for Life Sciences, Peking University, Beijing, P. R. China

**Author notes:** Correspondence: Yunlong Cao.

**Keywords:** Omicron breakthrough infection, Immune imprinting, Antibody responses, Epitope distribution

## Abstract

SARS-CoV-2 ancestral strain-induced immune imprinting poses great challenges to vaccine updates. Studies showed that repeated Omicron exposures could override immune imprinting induced by inactivated vaccines but not mRNA vaccines, a disparity yet to be understood. Here, we analyzed the underlying mechanism of immune imprinting alleviation in inactivated vaccine (CoronaVac) cohorts. We observed in CoronaVac-vaccinated individuals who experienced BA.5/BF.7 breakthrough infection (BTI), the proportion of Omicron-specific memory B cells (MBCs) substantially increased after an extended period post-Omicron BTI, with their antibodies displaying enhanced somatic hypermutation and neutralizing potency. Consequently, the neutralizing antibody epitope distribution encoded by MBCs post-BA.5/BF.7 BTI after prolonged maturation closely mirrors that in BA.5/BF.7-infected unvaccinated individuals. Together, these results indicate the activation and expansion of Omicron-specific naïve B cells generated by first-time Omicron exposure helped to alleviate CoronaVac-induced immune imprinting, and the absence of this process should have caused the persistent immune imprinting seen in mRNA vaccine recipients.

**Highlights:**

1. Longitudinal MBC profiling of CoronaVac-vaccinated individuals following BA.5 BTI
2. Omicron-specific MBC proportion rises greatly after extended period post-BA.5 BTI
3. Omicron-specific naive B cell maturation reduces ancestral strain immune imprinting

## Introduction

Immune imprinting refers to the tendency of the immune system to preferentially utilize memory from an initial viral encounter when dealing with subsequent infections with related but antigenically different strains^1^. This effect potentially compromises the effectiveness of responses elicited through subsequent vaccination or infections. Immune imprinting has been widely studied in influenza viruses, where individuals often exhibit a lifelong enhanced antibody response to the virus strains that were prevalent during their early years^2^.

Recently, this concept has gained renewed attention in the context of SARS-CoV-2. Among individuals who have been vaccinated with the SARS-CoV-2 ancestral strain, antibody responses to subsequent breakthrough infections (BTI) with variant strains are limited by immune imprinting^3–8^. Typically, this is characterized by the dominance of B cells that are cross-reactive to the ancestral strain, a low proportion of variant-specific B cells, and a significant proportion of antibodies targeting weakly neutralizing or non-neutralizing epitopes.

Previous studies reported minimum presence of Omicron-specific B cells and antibodies in individuals who experienced an Omicron BA.1 breakthrough infection after receiving mRNA vaccines targeting the ancestral strain^4–7^. Longitudinally, the proportion of BA.1-specific B cells after BA.1 BTI remains low, with enhanced responses to BA.1 largely attributable to the affinity maturation of cross-reactive memory B cells^9^. Even after two exposures to Omicron spikes, ancestral strain cross-reactive human memory B cells continued to dominate, rarely inducing Omicron-specific response^10^. Moreover, XBB.1.5 monovalent vaccines administered after an initial Omicron BTI failed to significantly boost Omicron-specific antibody responses^11^.

Conversely, in those immunized with inactivated vaccines, a modest proportion of Omicron-specific antibodies were observed 1-2 months following an Omicron breakthrough infection^3,8^. Remarkably, repeated exposures to the Omicron variant seem to significantly mitigate the effects of immune imprinting. This is evidenced by an increased prevalence of Omicron-specific antibodies, higher somatic hypermutation rates in these antibodies, and a shift towards more neutralizing epitopes. This observation leads us to hypothesize that the observed alleviation in immune imprinting could be a result of the expansion and maturation of Omicron-specific antibodies following an initial Omicron BTI.

It is crucial to decipher the differential behavior of immune imprinting between mRNA and inactivated vaccine recipients. In this study, we examined the immune imprinting alleviation mechanism of inactivated vaccine recipients. Specifically, we studied memory B cell (MBC) response 7 months after a BA.5/BF.7 BTI in individuals vaccinated with an inactivated vaccine. This analysis is compared to the MBC response observed shortly (1 month) after BA.5/BF.7 BTI, and the immune responses 8 months post-infection in individuals who have not been vaccinated but were infected with the BA.5/BF.7. Our analysis pays particular attention to the potential shifts in the epitope distribution of MBC encoded antibodies, aiming to elucidate the nuanced dynamics of immune responses to evolving SARS-CoV-2 variants. We demonstrated that, following a BA.5/BF.7 BTI in recipients of inactivated vaccines, there is significant expansion and maturation of *de novo* BA.5-specific naive B cells over an extended period. This substantially alters the epitope distribution within the long-term memory B cell antibody repertoire, resulting in an antibody epitope distribution that closely mirror those in unvaccinated infected cohorts. These results demonstrate that Omicron-specific naive B cell expansion and maturation reduces ancestral strain immune imprinting in inactivated vaccine recipients after an extended period following the first BTI.

## Results

### Expansion of BA.5-specific antibody encoded by memory B cell following an extended period post-BTI

To understand the long-term effects of immune imprinting elicited by inactivated vaccine, we analyzed immune responses in two cohorts who had experienced a BA.5/BF.7 infection more than half a year earlier. The first cohort consisted of individuals vaccinated with the inactivated vaccine based on the ancestral strain of SARS-CoV-2, who later experienced a BTI with the BA.5 or BF.7 variants; samples from this cohort were collected seven months post-infection (BA.5/BF.7 BTI (7 m)). The second cohort comprised unvaccinated individuals infected with the BA.5 or BF.7 variants, with samples collected eight months post-infection (BA.5/BF.7 inf. (8 m)). We also compared these cohorts with a previously reported cohort: individuals vaccinated with the WT SARS-CoV-2 inactivated vaccine who had a BTI with BA.5 or BF.7 variants, with samples collected one month post-infection to assess the initial immune response (BA.5/BF.7 BTI (1 m))^3,12^ (Figure 1A).

**Figure 1.**
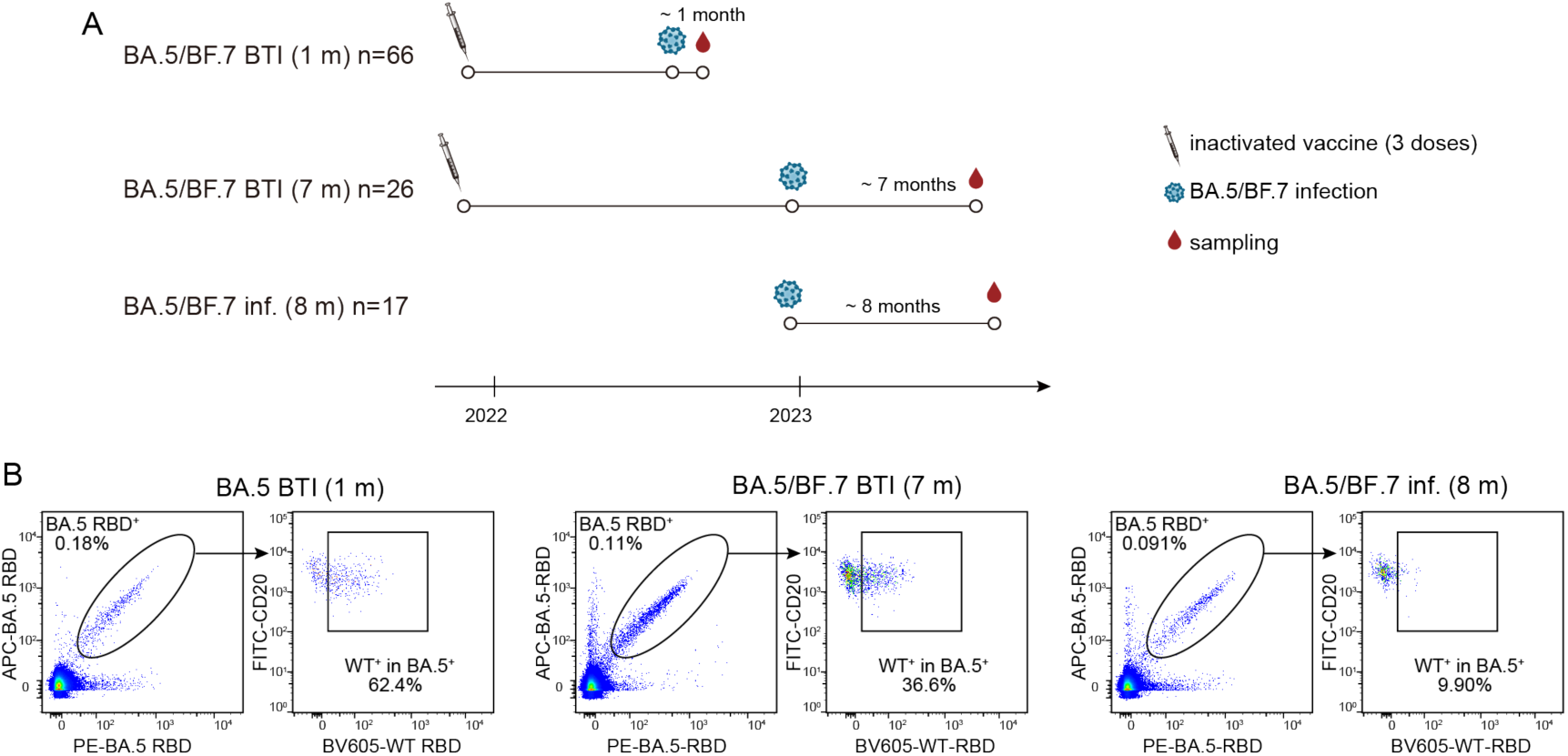
Memory B cell responses after an extended period post-BA.5 infection. (A) Timeline of vaccination, infection, and blood draws for 3 human cohorts, including short-term post-BA.5/BF.7 BTI sampling at 1 month (BA.5/BF.7 BTI (1 m)), prolonged post-BA.5/BF.7 BTI sampling at 7 months (BA.5/BF.7 BTI (7 m)), and prolonged post-BA.5/BF.7 infection sampling at 8 months (BA.5/BF.7 inf. (8 m)). (B) Flow cytometry analysis of pooled memory B cells from three cohorts including BA.5/BF.7 BTI (1 m), BA.5/BF.7 BTI (7 m), and BA.5/BF.7 inf. (8 m). BA.5 RBD APC and PE double-positive memory B cells (left panel) were analyzed for cross-reactivity with the wide-type (WT) RBD (right panel).

To assess the impact of immune imprinting on memory B cell responses, we analyzed B cells from the three cohorts using BA.5 and ancestral strain (wide-type, WT) receptor-binding domains (RBDs) as probes (Figure S1A). As a control, the BA.5/BF.7 inf. (8 m) cohort without previous vaccination history had the lowest cross-reactivity; in contrast, the majority of the B cells binding to BA.5 or BF.7 RBD also recognized the WT RBD in the BA.5 BTI (1 m) and BF.7 BTI (1 m) cohort as we previously reported^3^ (Figure 1B and Figure S1A). This cross-reactivity dropped significantly in the BA.5/BF.7 BTI (7 m) cohort, and a significant rise in proportion of BA.5-specific B cells was seen, which fell between the levels observed in the BA.5 BTI (1 m) and unvaccinated cohorts. This trend suggests the enduring expansion of BA.5-specific B cells after an extended period post-BTI, contrasting with the immediate aftermath of infection (Figure 1B).

To determine the exact proportions of specific and cross-reactive MBCs binding to BA.5 RBD, and to investigate whether these B cells undergo maturation along with expansion, BA.5 RBD-binding cells were isolated, and their paired heavy and light chain V(D)J were sequenced. Based on the sequences obtained, the corresponding antibodies were then synthesized *in vitro* as human IgG1monoclonal antibodies (mAbs). Among them, 189 mAbs were derived from the BA.5/BF.7 BTI (7 m) cohort, 195 mAbs from the BA.5/BF.7 inf. (8 m) cohort. Additionally, we included 628 mAbs previously identified from the BA.5/BF.7 BTI (1 m) cohort as controls to assess the maturation of the antibody repertoire over time post-BTI^3^. Using enzyme-linked immunosorbent assay (ELISA), we determined whether these antibodies were cross-reactive with WT strain or specific to BA.5 (Figure 2A). BA.5-specific antibodies accounted for 11% of all the antibodies in BA.5/BF.7 BTI (1 m), 50% in the BA.5/BF.7 BTI (7 m) cohort, and 70% in the BA.5/BF.7 inf. (8 m) cohorts. The outcomes of ELISA and flow cytometry analysis emphasized that BA.5-specific MBC expanded in proportion over an extended period following BA.5/BF.7 BTI in inactivated vaccine recipients.

**Figure 2.**
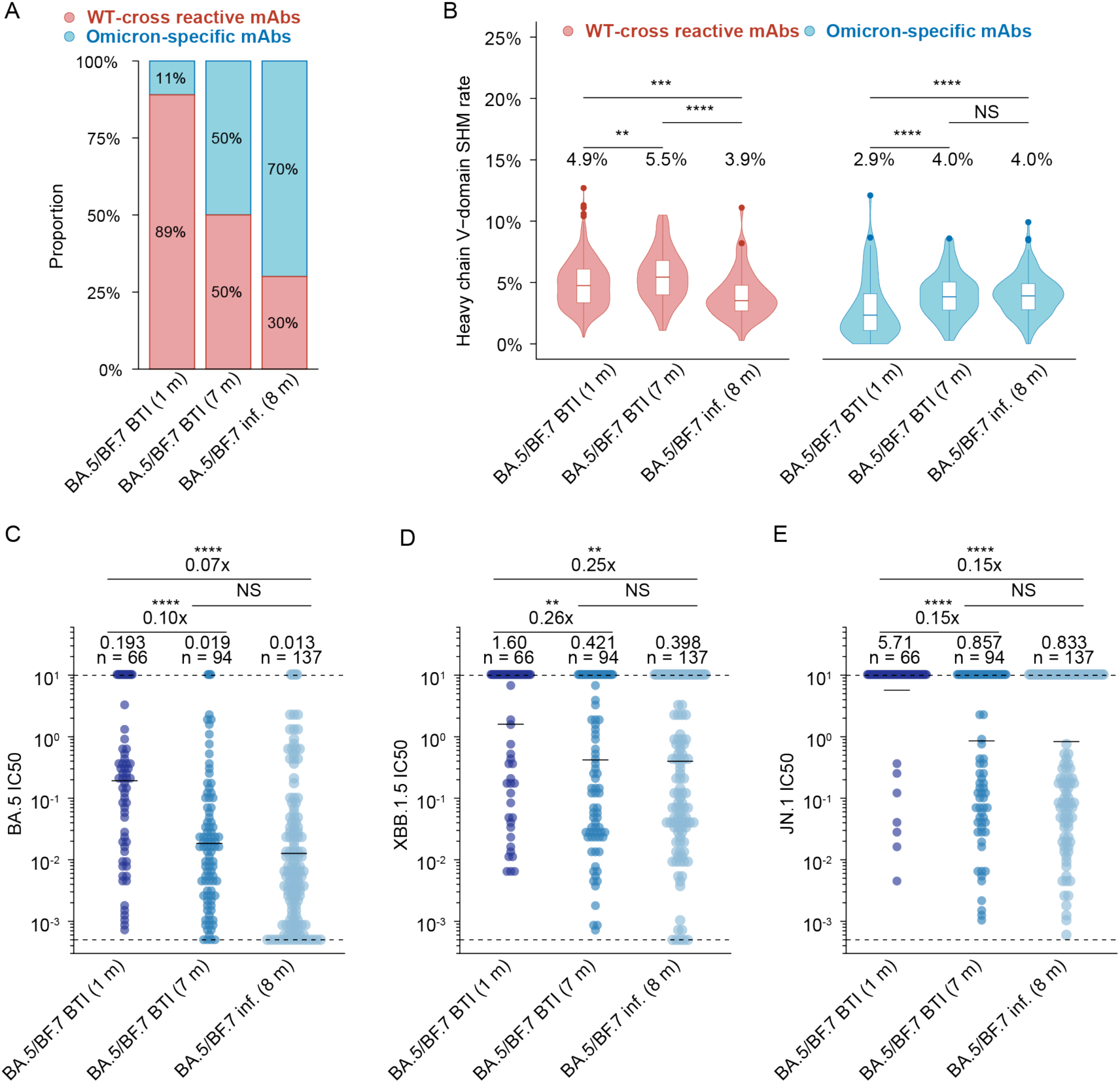
Expansion and maturation of BA.5 specific antibodies after prolonged time post-infection. (A) Proportions of WT cross-reactive and BA.5 specific antibodies from BA.5/BF.7 BTI (1 m), BA.5/BF.7 BTI (7 m), and BA.5/BF.7 inf. (8 m). The specificity of antibody binding was determined through ELISA. The antibodies were synthesized *in vitro* based on the sequences from BA.5-binding memory B cells. (B) The rate of somatic hypermutation in the heavy-chain variable domains of monoclonal antibodies (mAbs) derived from BA.5/BF.7 BTI (1 m), BA.5/BF.7 BTI (7 m), and BA.5/BF.7 inf. (8 m). Statistical analysis was conducted using two-tailed Wilcoxon rank-sum tests. The box plots represent the lower quartile, median, and upper quartile values, with whiskers extending to 1.5 times the interquartile range from the median. Violin plots illustrate the data’s distribution density. Details on the number and proportion of samples in each category are provided above the respective plots. (C) Half-maximal inhibitory concentration (IC50) of BA.5-specific mAbs from BA.5/BF.7 BTI (1 m), BA.5/BF.7 BTI (7 m), and BA.5/BF.7 inf. (8 m) against BA.5 (C), XBB (D), and JN.1 (D) pseudoviruses. The threshold for detection is marked by a dashed line. The geometric mean represented by a solid black bar. Annotations for geometric means, the variation in potency (fold changes), and antibody counts, were included. Statistical analysis was conducted using two-tailed Wilcoxon rank-sum tests. *P < 0.05, **P < 0.01, ***P < 0.001, ****P < 0.0001; NS, not significant (P > 0.05).

### Increased neutralization capacity of BA.5-specific antibodies due to maturation in the long-term response

Next, we examined the somatic hypermutation (SHM) rates and pseudovirus neutralizing activities of the MBC encoded antibodies across the cohorts. BA.5-specific MBC encoded antibodies of the two BA.5/BF.7 BTI cohorts demonstrated significantly lower SHM rates in both heavy and light chains than their cross-reactive counterparts, in alignment with their recent formation following BA.5/BF.7 BTI (Figure S2A-B). In contrast, antibodies from the individuals without ancestral strain vaccination exhibited similar SHM rates for both BA.5-specific and WT cross-reactive antibodies, indicating concurrent elicitation and comparable maturation periods (Figure S2A-B). Notably, by 7 months post-BTI, a significant increase in SHM rates of both heavy and light chains was observed for BA.5-specific antibodies relative to those observed at 1 month post-BTI, suggesting the maturation of BA.5-specific antibodies of MBC over time, and the SHM rates of BA.5-specific antibodies from the BA.5/BF.7 inf. (8 m) were similar to those from the BA.5/BF.7 BTI (7 m) given the comparable period of maturation, both considerably higher than those from BA.5/BF.7 BTI (1 m) (Figure 2B and Figure S2C).

We found the increase in SHM enhanced the neutralizing capacity of antibodies. BA.5-specific antibodies from both the BA.5/BF.7 BTI at 7 months and BA.5 infection at 8 months exhibited significantly greater neutralizing capabilities against BA.5, XBB.1.5 and JN.1 than those from the BA.5/BF.7 BTI at 1 month, suggesting the affinity maturation of antibodies encoded by MBC over time (Figure 2C-E). In comparison, WT cross-reactive antibodies from the BA.5/BF.7 BTI (7 m) and BA.5/BF.7 inf. (8 m) cohorts showed a modest but not significant improvement in neutralizing ability than those from shortly after infection, suggesting a comparatively minor role of the maturation process of cross-reactive antibodies in enhancing neutralization against Omicron (Figure S2D-F). These findings demonstrate that a prolonged period post-BTI significantly enhances the maturation of these BA.5-specific antibodies, making them more potent and broad compared to those observed shortly after BTI. This enhanced maturation explains why secondary exposure to Omicron can trigger highly potent serum neutralizing antibodies.

### Shifting of MBC-derived antibody epitope distribution after an extended period post-BA.5 BTI due to *de novo* BA.5-specific naïve B cell maturation

We then analyzed the differences in antibody epitope distribution and compared them between vaccinated and unvaccinated cohorts after a prolonged period following BA.5/BF.7 infection. The epitopes of antibodies were determined by performing yeast-display-based high throughput deep mutational scanning (DMS) on the BA.5 RBD, as previously described^12,13^. Accordingly, twelve major epitope groups were identified by applying graph-based unsupervised clustering to the escape scores of each antibody across BA.5 RBD sites. For visual representation, we displayed the epitope grouping of antibodies using uniform manifold approximation and projection (UMAP) (Figure S3A-G), and representative antibodies from each epitope group were shown in a structural complex with the RBD (Figure 3A).

**Figure 3.**
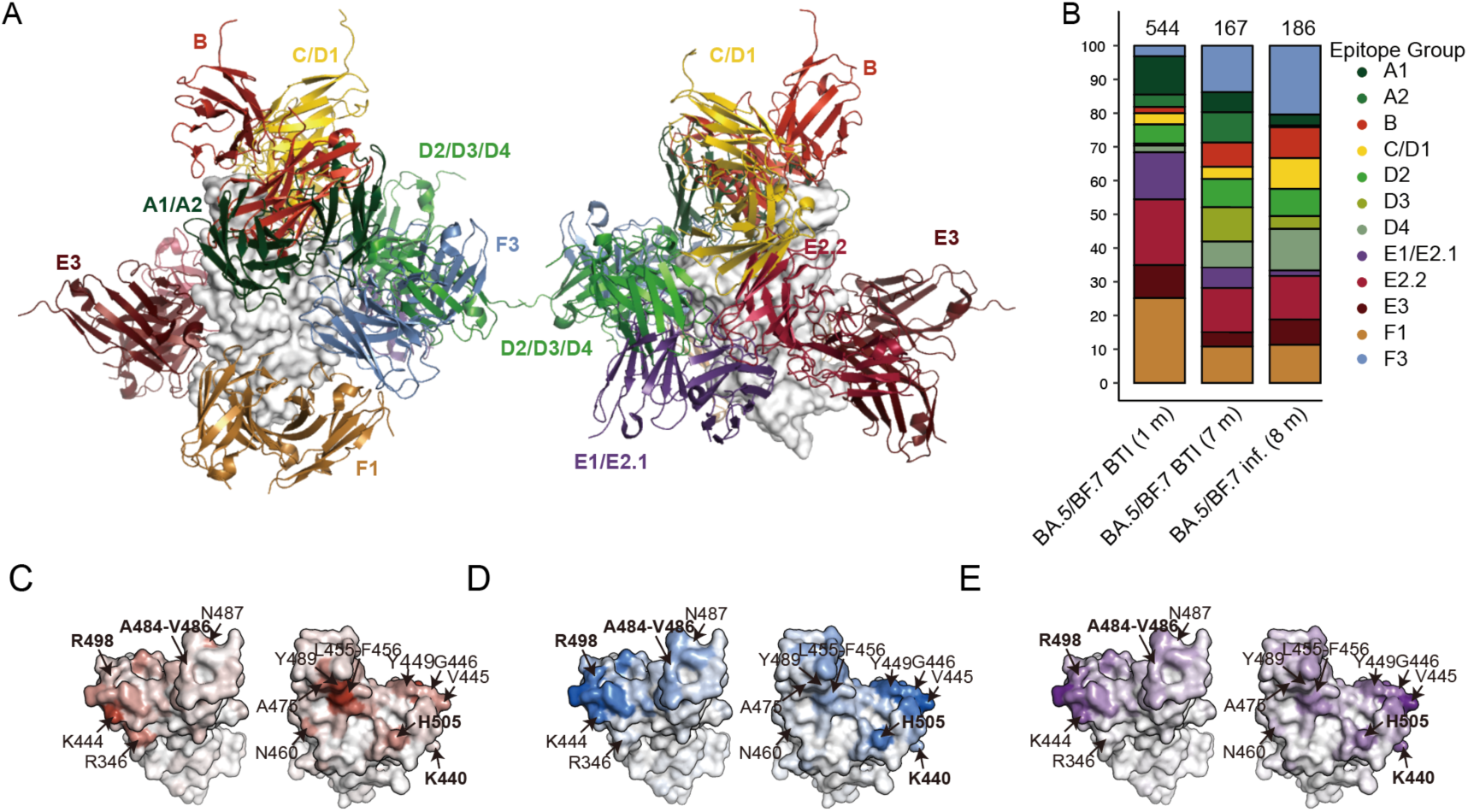
Epitope distribution of antibodies after prolonged time post-infection. (A) Illustration of antibody epitope groups of A1-F3. Structures of RBD and representative antibody of each group are shown. (B) Distribution of monoclonal antibodies among different epitope groups is shown for BA.5/BF.7 BTI (1 m), BA.5/BF.7 BTI (7 m), and BA.5/BF.7 inf. (8 m). Number of mAbs are shown above the column. (C-E) Normalized average DMS escape scores, weighted by IC50 against BA.5 using DMS profiles of monoclonal antibodies from BA.5/BF.7 BTI (1 m) (C), BA.5/BF.7 BTI (7 m) (D), and BA.5/BF.7 inf. (8 m) (E), are indicated on the structure model of SARS-CoV-2 BA.5 RBD (PDB: <u>7XNS</u>). Residues with change in immune pressure are labelled. Residues with immunogenicity change in BA.5 compared to ancestral strain are labelled in bold.

Importantly, there was a shift in epitope distribution over time following the BA.5/BF.7 BTI. The epitope distribution of antibodies at 7 months post-BTI was notably different from that at 1 month post-BTI, yet closely aligned with that seen in individuals 8 months post-BA.5/BF.7 infection (Figure 3B). Compared to BA.5/BF.7 (1 m), there is an increase in antibodies targeting neutralizing Omicron-specific epitope groups B, D2, D3, D4, and F3, and a decrease in antibodies targeting weak neutralizing cross-reactive epitope groups E2.2, E3, and F1 (Figure S3C-F).

To further explore the differences of antibodies elicited by BA.5/BF.7 BTI at 7 months compared to those from the BA.5/BF.7 BTI at 1 month, we determined the cumulative escaping score for antibodies from each cohort and adjusted it based on their neutralizing activity against BA.5, which is referred to as immune pressure distribution (Figure S3H). Immune pressure sites are highlighted on the BA.5 RBD illustration, indicating that antibodies targeting these sites are more numerous and potent (Figure 3C-E). Indeed, the immune pressure sites for antibodies at 7 months post-BTI significantly differed from those at 1 month post-BTI, and some of these sites have undergone mutations in BA.5 compared to ancestral strain, emphasizing the changes in immunogenicity. Specifically, mutations at sites such as 440, 484, 486, 498, and 505 could more readily evade antibodies from the BA.5/BF.7 BTI at 7 months than those from the 1 month post-BTI. These sites have undergone mutations that alter immunogenicity in BA.5, such as N440K (a change from polar uncharged to positively charged side chains), E484A (from negatively charged to hydrophobic), Q489R (from polar uncharged to positively charged), and Y505H (from hydrophobic to positively charged). Furthermore, the mutation sites escaping antibodies of the BA.5/BF.7 BTI at 7 months closely resemble the sites identified in the BA.5/BF.7 infection at 8 months (Figure 3C). These observations again emphasize that the antibodies targeting these mutated sites were specifically generated and matured in response to the BA.5 exposure. Moreover, when analyzing the heavy chain V gene usage of these antibodies, it was observed that the usage of BA.5-specific antibodies at 7 months post-BTI was similar to that at 1 month post-BTI and in BA.5/BF.7 infections, but distinct from the cross-reactive antibodies at 1 month post-BTI (Figure 3D). These results suggest the shift in epitope distribution of antibodies encoded by MBC in BA.5/BF.7 BTI (7 m) compared to BA.5/BF.7 BTI (1 m) is caused by the maturation of *de novo* generated BA.5-specific naïve B into MBC.

## Discussion

Our results indicate that in recipients of inactivated vaccines, Omicron-specific naïve B cells significantly expand, mature, and evolve into memory B cells over an extended period following an Omicron breakthrough infection. The induction, growth, and maturation of Omicron-specific naïve B cells, culminating in their conversion to MBCs following first Omicron BTI, play a key role in reducing immune imprinting effects during later encounters with the variant. These Omicron specific MBC can be efficiently reactivated upon a second exposure to Omicron, leading to the production of high levels of neutralizing antibodies^12^.The absence of such inducement and expansion of Omicron-specific antibodies in individuals who received ancestral strain mRNA vaccines may explain the sustained immune imprinting even upon re-exposure to Omicron. This may be attributed to the high immunogenicity of mRNA vaccines^14^, which trigger a robust immune response, leading to an abundance of potent memory B cells and antibodies. This extensive pre-existing immunity may impede the recruitment of naïve B cells to variant strains during Omicron exposures^15–17^. It is crucial to unravel the reasons why ancestral strain mRNA vaccine recipients lack Omicron-specific antibodies after BTI, because this insight helps to develop vaccines that foster variant-specific B cell responses, effectively overcoming immune imprinting and enhancing Omicron-targeting antibodies.

### Limitation of the study

One limitation of our study was the inability to collect samples from the same individuals at both earlier and later time points. Nevertheless, we believe that the data averaged across different individuals may still reflect general trends. We also could not obtain samples from unvaccinated individuals shortly after BA.5/BF.7 infection. While the magnitude of antibody responses may vary between short-term and long-term observations, we propose that the nature of the antibodies, including their specificity and epitopes, likely remains consistent over time in these individuals.

## Method

### Key resource table

KEY RESOURCES TABLE
| REAGENT or RESOURCE | SOURCE | IDENTIFIER |
| --- | --- | --- |
| Antibodies |  |  |
| FITC anti-human CD20 Antibody | BioLegend | 302304 |
| Brilliant Violet 421™ anti-human CD27 Antibody | BioLegend | 302824 |
| PE/Cyanine7 anti-human IgM Antibody | BioLegend | 314532 |
| PE/Cyanine7 anti-human IgD Antibody | BioLegend | 348210 |
| Brilliant Violet 421™ anti-human CD27 Antibody | BioLegend | 302824 |
| PE/Cyanine7 anti-human IgM Antibody | BioLegend | 314532 |
| PE/Cyanine7 anti-human IgD Antibody | BioLegend | 348210 |
| Peroxidase AffiniPure™ Goat Anti-Human IgG (H+L) | JACKSON | Cat#109-035-003<br>RRID: AB_2337577 |
| HG1K | Sino Biological | Cat#HG1K |
| Chemicals, Peptides, and Recombinant Proteins |  |  |
| BA.5 biotinylated RBD | Sino Biological | Cat# 40592-V49H9-B |
| PE-streptavidin | BioLegend | Cat#405204 |
| PC-streptavidin | BioLegend | Cat#405207 |
| BV605-streptavidin | BioLegend | Cat#405229 |
| WT biotinylated RBD | Sino Biological | Cat#40592-V27H-B |
| 7-AAD | Invitrogen | Cat#00-6993-50 |
| ELISA coating buffer | Solarbio | Cat#C1055 |
| Recombinant Human ACE2 Protein (His Tag), Amine-Labeled Biotinylated | Sino Biological | Cat#10108-H08H-B |
| Critical Commercial Assays |  |  |
| Human CD19 Positive Selection Kit II | STEMCELL | Cat#17854 |
| Chromium Next GEM Single Cell 5' Library & Gel Bead Kit v1.1, 16 rxns | 10X Genomics | Cat#PN-1000165 |
| Chromium Single Cell V(D)J Enrichment Kit, Human B Cell, 96 rxns | 10X Genomics | Cat#PN-1000016 |
| Chromium Next GEM Chip G Single Cell Kit, 16 rxns | 10X Genomics | Cat#PN-1000127 |
| Single Index Kit T Set A, 96 rxns | 10X Genomics | Cat#PN-1000213 |
| ClonExpress II One Step Cloning Kit | Vazyme | Cat#C112 |
| EndoFree Plasmid Midi Kit | CWBIO | Cat#CW2105 |
| Bright-Lite Luciferase Assay System | Vazyme | Cat#DD1204 |
| Deposited Data |  |  |
| Structure model of BA.5 RBD | Cao et al. 2022 | PDB: 7XNS |
| DMS profiles of SARS-CoV-2 RBD-targeting mAbs elicited by BA.5/BF.7 breakthrough infections | Cao et al. 2023 | N/A |
| Experimental Models: Cell Lines |  |  |
| Expi293F human cells | Thermo Fisher | Cat#A14527 |
| HuH-7 cells | JCRB | Cat#JCRB0403 |
| Vero cells | ATCC | Cat#CCL-81 |
| 293T cells | ATCC | Cat#CRL-3216 |
| Experimental Models: Organisms/Strains |  |  |
| *Saccharomyces cerevisiae* strain EBY100 | ATCC | Cat#MYA-4941 |
| Trelief® 5α Chemically Competent Cell | Tsingke | Cat#TSC-C01-96 |
| Pseudotyped ΔG-luciferase (G*ΔG-luciferase) rVSV | Kerafast | Cat#EH1020-PM |
| Oligonucleotides |  |  |
| NEBNext® Multiplex Oligos for Illumina® (96 Index Primers) | New England Biolabs | Cat#E6609L |
| Recombinant DNA |  |  |
| pETcon_SARS-CoV-2_RBD | Addgene | Cat#166782; RRID:Addgene_166782 |
| pCMV3 | Sino Biological | Cat#CV011 |
| Software and Algorithms |  |  |
| Cell Ranger (v6.1.1) | 10x Genomics | https://support.10xgenomics.com/single-cell-gene-expression/software/pipelines/latest/installation |
| IgBlast (v1.17.1) | NCBI | https://ncbi.github.io/igblast/ |
| Change-O (v1.2.0) | Gupta NT et al., 2015 | *doi: 10.1093/bioinformatics/btv359* |
| Summit (v6.0) | Beckman Coulter | https://www.beckmancoulter.com/ |
| FlowJo (v10.8) | BD Biosciences | https://www.flowjo.com |
| PRISM (v 9.0.1) | Graphpad | https://www.graphpad.com/updates/prism-901-release-notes |
| PyMOL (v2.6.0a0) | Schrödinger | https://github.com/schrodinger/pymol-open-source |
| ggplot2 (v3.3.3) | Wickham et al. 2009 | https://ggplot2.tidyverse.org/ |
| dms_variants (v1.5.0) | J. Bloom lab | https://github.com/jbloomlab/dms_variants/ |
| umap-learn (v0.5.2) | McInnes et al. arXiv 2018 | https://github.com/lmcinnes/umap |
| leidenalg (v0.10.2) | Traag et al. 2018 | https://github.com/vtraag/leidenalg |
| scipy (v1.7.0) | Virtanen et al. 2020 | https://scipy.org/ |
| Other |  |  |
| AmMag™ Protein A Magnetic Beads | Genscript | Cat#L00695 |
| Dynabeads™ Protein A | Thermo Fisher | Cat#10008D |
| Pierce™ Anti-c-Myc Magnetic Beads | Thermo Fisher | Cat#88843 |
| Dynabeads® Biotin Binder | Thermo Fisher | Cat#11047 |

## Experimental model and study participant details

### Cell lines

The 293 T cell line (ATCC, CRL-3216) is a female cell line. Huh-7 cell line (JCRB, 0403) is a male cell line. Expi293F (Thermo Fisher, A14527) human cells are from a female cell line. Cells were maintained at a temperature of 37°C. Adherent cells were cultivated in an atmosphere of 5% CO_2_ and suspension cells were cultivated at 8% CO_2_ with 130 rpm of agitation.

### Sample donors and collection

Blood samples were collected after informed consent from vaccinated or unvaccinated individuals who had BA.5/BF.7 breakthrough infection history. Study protocols received approval from Beijing Ditan Hospital, Capital Medical University (Ethics committee archiving no. LL-2021-024-02) and the Ethics Committee of Tianjin First Central Hospital (Ethics committee archiving no. 2022N045KY). Detailed information is provided in Supplementary Table 1.

## Method details

### PBMC and plasma isolation

Whole blood samples were diluted at a ratio of 1:1 using 2% FBS PBS, and then processed using a Ficoll density gradient centrifugation technique for the separation of plasma and peripheral blood mononuclear cells (PBMCs). After centrifugation, plasma was retrieved from the top layer. PBMCs were collected from the boundary layer, followed by additional centrifugation, lysis of red blood cells using 1× RBC Lysis Buffer and washing steps. PBMCs were preserved in FBS enhanced with 10% DMSO and stored in liquid nitrogen. All PBMC samples were transported using dry ice. For subsequent applications, the cryopreserved PBMCs were gently thawed in a solution of PBS with 1 mM EDTA and 2% FBS.

### BCR sequencing and analysis

CD19+ B cells were separated from PBMCs using the EasySep Human CD19 Positive Selection Kit II. After B cell isolation, for each sample containing 1 × 10^6 B cells in 100 μL of 2% FBS PBS, a mixture of antibodies was added: 3 μl FITC anti-human CD20 antibody, 3.5 μl Brilliant Violet 421 anti-human CD27 antibody, 2 μl PE/Cyanine7 anti-human IgD antibody, and 2 μl PE/Cyanine7 anti-human IgM antibody. Additionally, 0.013 μg of RBD tagged with fluorophores was added. Following a 30-minute incubation and two wash cycles, 5 μl of 7-AAD was incorporated to discern live cells.

Cells devoid of 7-AAD, IgM, and IgD staining but positive for CD20, CD27, and BA.5 RBD were sorted using the MoFlo Astrios EQ Cell Sorter (Beckman Coulter). The data from FACS were gathered using Summit 6.0 and analyzed with FlowJo software version 10.8.

The isolated RBD-binding B cells were prepared for sequencing using the Chromium Next GEM Single Cell V(D)J Reagent Kits v1.1, following the guidelines provided by the manufacturer (10X Genomics, CG000208). In brief, the RBD-binding B cells were reencapsulated into gel beads-in-emulsion (GEMs) using the 10X Chromium controller. The GEMs underwent reverse transcription. Then the products were further purified. Products were subject to preamplification and were subsequently purified with SPRIselect Reagent Kit. The paired V(D)J sequences were amplified with 10X BCR primers, followed by sequencing library preparation. Sequencing of these libraries was then performed on the Novaseq 6000 platform.

V(D)J sequencing data from 10X Genomics were compiled into BCR contigs with Cell Ranger according to GRCh38 BCR reference sequence. Criteria were set to include only those BCR contigs that were productive, as well as cells that exhibited one single heavy chain and one single light chain. The germline V(D)J genes were annotated and somatic hypermutation sites within the variable regions of the BCR sequences were detected using IgBlast (v1.17.1) and Change-O (v1.2.0).

### Antibody synthesis

Heavy and light chain genes were optimized and synthesized by GenScript, inserted separately into plasmids pCMV3-CH and pCMV3-CL or pCMV3-CK with ClonExpressII One Step Cloning Kit. The recombinant products were transformed into DH5α competent cells and plated on LB agar containing ampicillin. After overnight cultures, positive single colonies were picked for PCR identification. Correct clones were selected, cultured for expansion, and plasmid extraction was performed using the EndoFree Plasmid Midi Kit. Plasmids were then co-introduced into Expi293F cells via transfection mediated by polyethylenimine. Post-transfection, the cells were cultured for 6-10 days. The supernatant was harvested and then processed for purification using AmMag™ protein A magnetic beads.

### VSV pseudotyped virus production

Vesicular stomatitis virus (VSV) was pseudotyped to display Spike protein of SARS-CoV-2, incorporating mutaions of D614G, BA.5, XBB.1.5, or JN.1 respectively, following a previously described protocol^18^. Briefly, the 293T cells were introduced to plasmids encoding the codon-optimized variant spike gene, followed by infection with pseudotyped ΔG-luciferase (G*ΔG-luciferase) rVSV. After cell culture, the supernatants containing pseudotyped VSV were harvested, centrifuged, filtered, aliquoted, and frozen at −80 °C.

### Pseudotyped VSV neutralization

Monoclonal antibodies or plasma samples were sequentially diluted within DMEM medium and then mixted with pseudotyped VSV samples in 96-well plates, maintained at 37°C 5% CO_2_ for one hour. Subsequently, digested Huh-7 cells were added to the wells and incubated for 24 hours. Afterwards, half of the medium was removed, and Bright-Lite Luciferase Assay System was introduced into the wells for reaction in absence of light. The luminescence was measured with a microplate spectrophotometer. The half-maximal inhibitory concentration (IC50) or half-maximal neutralizing titers were calculated employing a four-parameter logistic regression analysis, facilitated by the PRISM.

### ELISA

ELISA assays were performed by initially coating ELISA plates with RBD in ELISA coating buffer, and incubating overnight at 4°C. The plates were then undergone washing and blocking processes. Subsequently, antibodies were dispensed into each well and left to incubate at ambient temperature for 30 minutes. After incubation, the plates were washed again and subsequently incubated with Peroxidase-conjugated AffiniPure goat anti-human IgG (H+L) for 30 minutes at room temperature. The reaction was developed with tetramethylbenzidine (TMB), and halted with H2SO4. The absorbance was measured at 450 nm using a microplate reader. HG1K, a human IgG1 antibody against H7N9, served as the negative control.

### High-throughput antibody-escape mutation profiling

High-throughput antibody-escape mutation profiles were conducted using the previously constructed duplicated deep mutational scanning (DMS) libraries based on receptor binding domain (RBD) of BA.5 variant^12^. As previously described^3,8,13^, the successfully expressed and properly folded RBD variants in DMS libraries were first enriched by magnetic beads, and then further enlarged and induced for downstream mutation escape profiling. To eliminate the antibody binding RBD variants as much as possible, two rounds of negative cell sorting were performed by incubating the above enriched libraries with antibody conjugated Protein A magnetic beads. These antibody escapers were proceeded to positive cell sorting using the anti-c-Myc tag magnetic beads to further eliminate yeast cells without expressed RBD variants. The finally obtained yeast population was expanded by overnight growth in SD-CAA media and lysed for plasmid extraction. The next generation sequencing libraries were prepared by PCR amplifying the region spanning the N26 barcode on the plasmids and submitted to Illumina NextSeq 550 or MGI Tech MGISEQ-2000 platform.

Raw NGS data of barcodes were processed and the escape scores of each antibody and mutation were calculated by custom scripts similar to the methods in our previous report. In brief, we calculated the ratio of fraction of barcodes corresponding to each variant between the antibody-selected library and the reference library. The ratios of fractions were scaled to 0-1 range, and an epistasis model was fitted using dms_variants (v1.5.0) to get the escape scores of each single substitution. For clustering analysis, site escape scores (the total escape scores on a residue over different substitutions) of each antibody are first normalized to a summation of one and considered as a distribution over RBD residues. Dissimilarities of each pair of antibodies are defined as the Jessen-Shannon divergence of normalized escape scores, and calculated using scipy (v1.7.0). Then, igraph (v0.9.6) was used to build a 12-nearest-neighbor graph and performed leiden clustering (leidenalg v0.10.2) to assign a cluster to each antibody. The epitope group of each cluster is manually annotated based on the featured sites of each cohort to ensure the consistency with the definition of our previously published SARS-CoV-2 RBD DMS datasets. UMAP was performed based on the constructed k-nearest-neighbor graph using umap-learn module (v0.5.2) to project the antibody epitope profiles onto a 2D space for visualization. Within this antibody clustering dataset, antibody counts of 702, 167, and 186 corresponded to samples from BA.5/BF.7 BTI at 1 month, BA.5/BF.7 BTI at 7 month, and BA.5/BF.7 infections at 8 months, respectively (Figure S3B). We also incorporated antibodies from earlier collections that were isolated from diverse immunological backgrounds to better elucidate the characteristics of antibodies across different epitopes. Illustration was performed by R package ggplot2 (v3.3.3), and pymol (v2.6.0a0).

## Acknowledgments

This study was financially supported by the Ministry of Science and Technology of China and Changping Laboratory (2021A0201 and 2021D0102 to Y.C.), and National Natural Science Foundation of China (32222030 to Y.C.).

## Author contributions

Y.C. conceived and supervised the study. A.Y. managed the study. A.Y, W.S., F.J., and Y.C. wrote the manuscript with inputs from all authors. A.Y., W.S., R.A., Y.W., J.W(Changping Laboratory), and F.S. performed B cell sorting, single-cell V(D)J sequencing, and antibody sequence analyses, and antibody expression. J.W. (BIOPIC), F.J., and H.S. performed and analysed the DMS data. Y.Y. and Youchun Wang constructed the pseudotyped virus. P.W., L.Y., and F.S. performed the pseudotyped virus neutralization assays and ELISA and SPR. W.S. and A.Y. analysed the neutralization data. X.C., Y.X., R.J., and Z.S. recruited the SARS-CoV-2 BA.5/BF.7 convalescents.

## Figures

**Figure S1.**
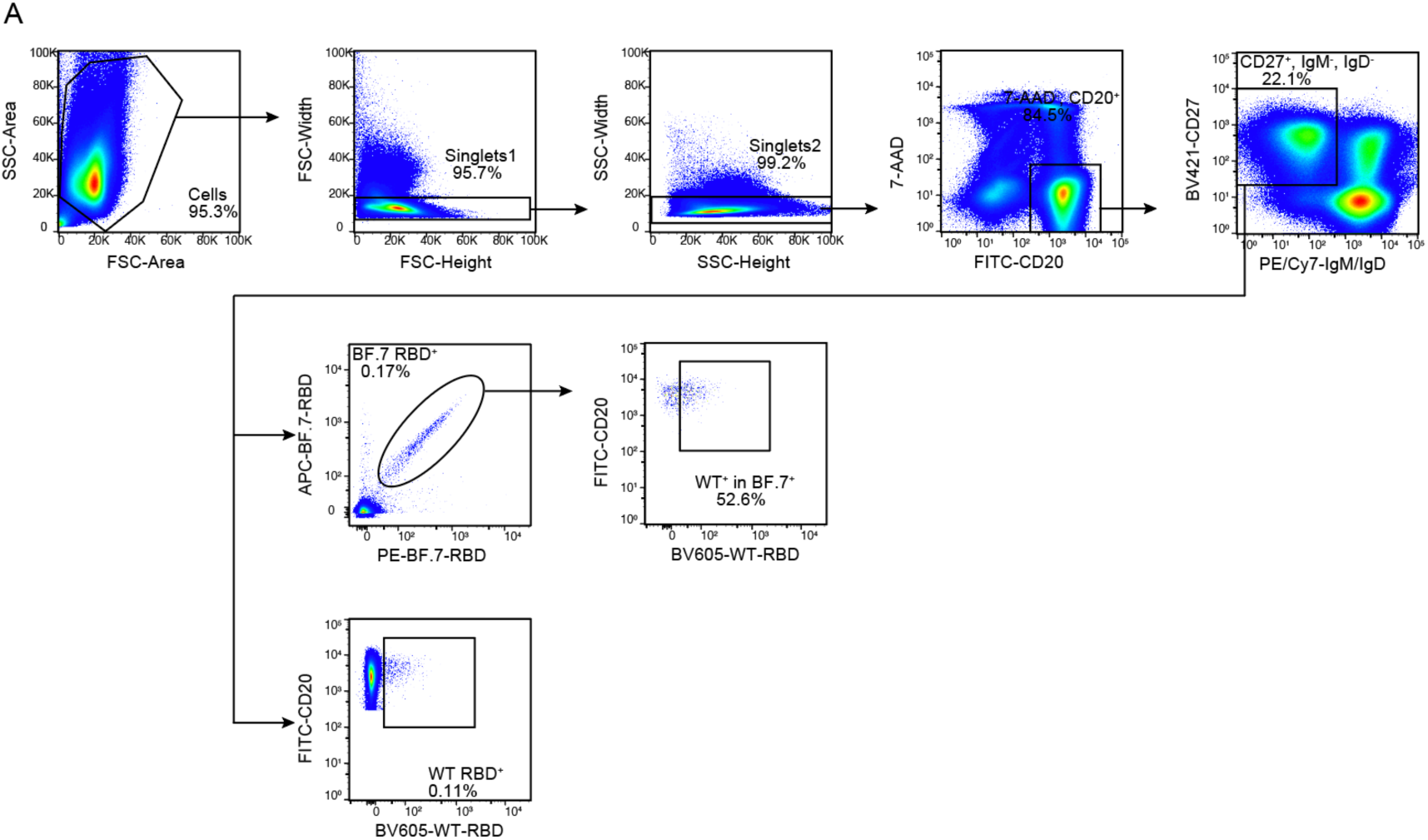
related to Figure 1 Memory B cell response after BA.5/BF.7 infection. (A) Gating strategy for assessing the cross-reactivity of BA.5 or BF.7 RBD ^+^memory B cells with the WT RBD. Individuals with BF.7 BTI (1 m) were shown as an example.

**Figure S2.**
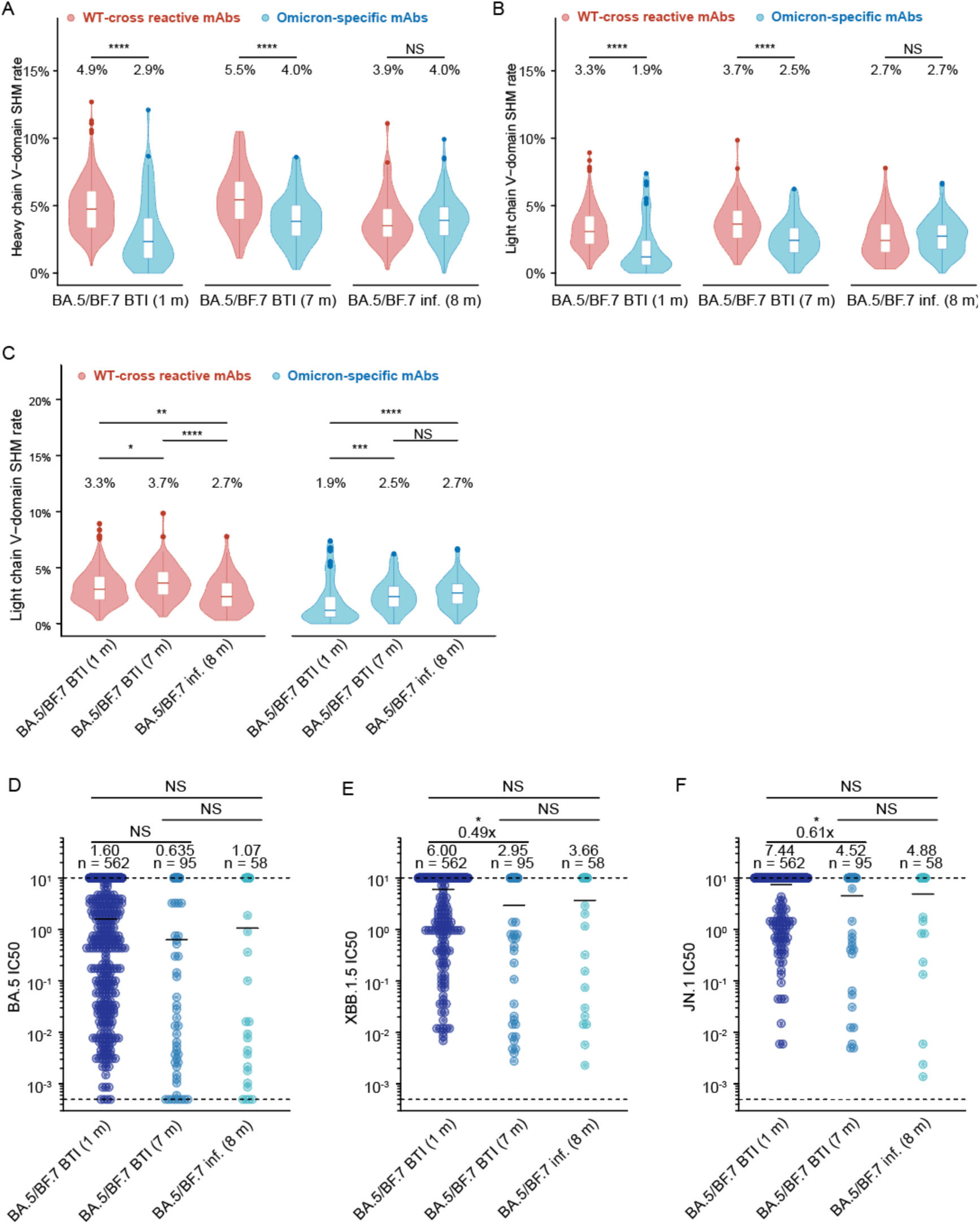
related to Figure 2 Maturation of antibodies after BA.5 infection. (A-B) Comparison of somatic hypermutation of Omicron-specific and WT-cross reactive in the heavy-chain variable domains (A) and light-chain vatiable domains (B) of monoclonal antibodies (mAbs) derived from BA.5/BF.7 BTI (1 m), BA.5/BF.7 BTI (7 m), and BA.5/BF.7 inf. (8 m). (C) The rate of somatic hypermutation in the light-chain variable domains of monoclonal antibodies (mAbs) derived from BA.5/BF.7 BTI (1 m), BA.5/BF.7 BTI (7 m), and BA.5/BF.7 inf. (8 m). Statistical analysis for (A-C) was conducted using two-tailed Wilcoxon rank-sum tests. The box plots represent the lower quartile, median, and upper quartile values, with whiskers extending to 1.5 times the interquartile range from the median. Violin plots illustrate the data’s distribution density. Details on the number and proportion of samples in each category are provided above the respective plots. (D-F) Half-maximal inhibitory concentration (IC50)of WT cross-reactive mAbs from BA.5/BF.7 BTI (1 m), BA.5/BF.7 BTI (7 m), and BA.5/BF.7 inf. (8 m) against BA.5 (D), XBB (E), and JN.1 (F) pseudoviruses. The threshold for detection is marked by a dashed line. The geometric mean represented by a solid black bar. Annotations for geometric means, the variation in potency (fold changes), and antibody counts, were included. Statistical analysis was conducted using two-tailed Wilcoxon rank-sum tests.

**Figure S3.**
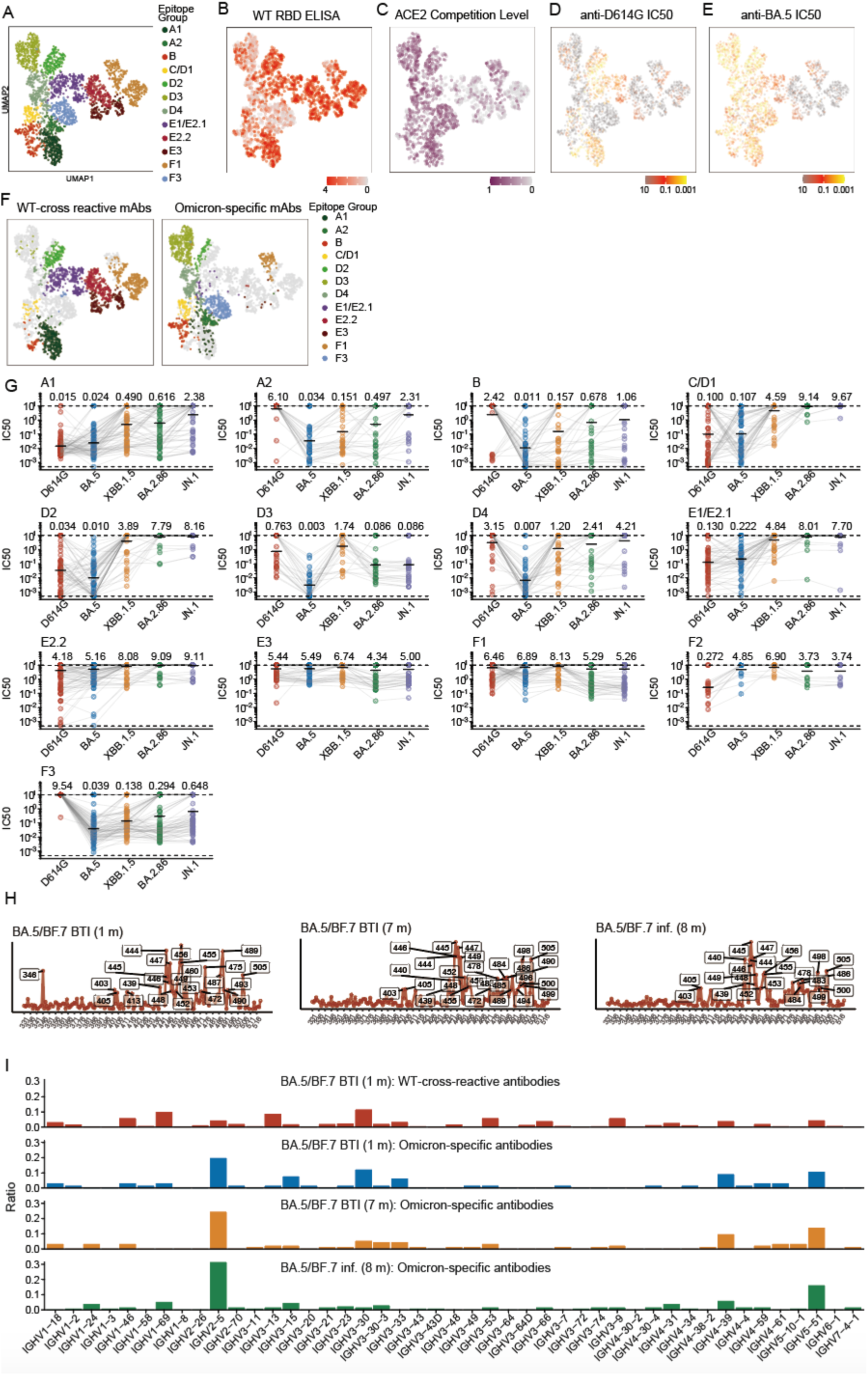
related to Figure 3 Epitope distribution and characterization of monoclonal antibodies. (A) UMAP embedding of epitope groups of monoclonal antibodies binding BA.5 RBD (n = 2294). (B) WT binding affinity of monoclonal antibodies from BA.5/BF.7 BTI (1 m), BA.5/BF.7 BTI (7 m), BA.5/BF.7 inf. (8 m), determined by ELISA, denoted as OD450 value, are projected onto the UMAP embedding space (n = 2051). (C) ACE2 competition monoclonal antibodies from BA.5/BF.7 BTI (1 m), BA.5/BF.7 BTI (7 m), BA.5/BF.7 inf. (8 m), determined by ELISA, are projected onto the UMAP embedding space (n = 1662). (D-E) Neutralization activities against D614G and BA.5 spike-pseudovirus, denoted as IC50 values, are projected onto the UMAP embedding space (n = 2294 and n =2292). (F) UMAP embedding of epitope groups of WT-cross-reactive mAbs and Omicron-specific mAbs. (G) Pseudovirus-neutralization activities of monoclonal antibodies in each epitope groups are shown against SARS-CoV-2 D614G, BA.5, XBB.1.5, BA.2.86, and JN.1. Geometric mean IC50 values are displayed as bars and indicated above each group of data points. A1 (n = 236); A2 (n = 100); B (n = 139); C/D1 (n = 84); D2 (n = 166); D3 (n = 272); D4 (n = 210); E1/E2.1 (n = 245); E2.2 (n = 208); E3 (n = 124); F1 (n = 307); F3 (n = 203) (H) Immune pressure of each cohort determined by the cumulative escaping score for antibodies from each cohort and adjusted it based on their neutralizing activity against BA.5. (G) Heavy chain gene usage distribution of WT-cross-reactive antibodies and Omicron specific antibodies from BA.5/BF.7 BTI(1 m) and Omicron specific antibodies from BA.5/BF.7 BTI(7 m) and BA.5/BF.7 inf. (8 m). n= 562, 66, 94,137, respectively.

## Reference

1. Francis, T. (1960). On the Doctrine of Original Antigenic Sin. Proceedings of the American Philosophical Society 104, 572–578.

2. Lessler, J., Riley, S., Read, J.M., Wang, S., Zhu, H., Smith, G.J., Guan, Y., Jiang, C.Q., and Cummings, D.A. (2012). Evidence for antigenic seniority in influenza A (H3N2) antibody responses in southern China. PLoS Pathog 8, e1002802. 10.1371/journal.ppat.1002802.

3. Cao, Y., Jian, F., Wang, J., Yu, Y., Song, W., Yisimayi, A., Wang, J., An, R., Chen, X., Zhang, N., et al. (2023). Imprinted SARS-CoV-2 humoral immunity induces convergent Omicron RBD evolution. Nature 614, 521–529. 10.1038/s41586-022-05644-7.

4. Alsoussi, W.B., Malladi, S.K., Zhou, J.Q., Liu, Z., Ying, B., Kim, W., Schmitz, A.J., Lei, T., Horvath, S.C., Sturtz, A.J., et al. (2023). SARS-CoV-2 Omicron boosting induces de novo B cell response in humans. Nature 617, 592–598. 10.1038/s41586-023-06025-4.

5. Kaku, C.I., Bergeron, A.J., Ahlm, C., Normark, J., Sakharkar, M., Forsell, M.N.E., and Walker, L.M. (2022). Recall of preexisting cross-reactive B cell memory after Omicron BA.1 breakthrough infection. Sci Immunol 7, eabq3511. 10.1126/sciimmunol.abq3511.

6. Park, Y.J., Pinto, D., Walls, A.C., Liu, Z., De Marco, A., Benigni, F., Zatta, F., Silacci-Fregni, C., Bassi, J., Sprouse, K.R., et al. (2022). Imprinted antibody responses against SARS-CoV-2 Omicron sublineages. Science 378, 619–627. 10.1126/science.adc9127.

7. Quandt, J., Muik, A., Salisch, N., Lui, B.G., Lutz, S., Kruger, K., Wallisch, A.K., Adams-Quack, P., Bacher, M., Finlayson, A., et al. (2022). Omicron BA.1 breakthrough infection drives cross-variant neutralization and memory B cell formation against conserved epitopes. Sci Immunol 7, eabq2427. 10.1126/sciimmunol.abq2427.

8. Cao, Y., Yisimayi, A., Jian, F., Song, W., Xiao, T., Wang, L., Du, S., Wang, J., Li, Q., Chen, X., et al. (2022). BA.2.12.1, BA.4 and BA.5 escape antibodies elicited by Omicron infection. Nature 608, 593–602. 10.1038/s41586-022-04980-y.

9. Kaku, C.I., Starr, T.N., Zhou, P., Dugan, H.L., Khalife, P., Song, G., Champney, E.R., Mielcarz, D.W., Geoghegan, J.C., Burton, D.R., et al. (2023). Evolution of antibody immunity following Omicron BA.1 breakthrough infection. Nat Commun 14, 2751. 10.1038/s41467-023-38345-4.

10. Addetia, A., Piccoli, L., Case, J.B., Park, Y.J., Beltramello, M., Guarino, B., Dang, H., de Melo, G.D., Pinto, D., Sprouse, K., et al. (2023). Neutralization, effector function and immune imprinting of Omicron variants. Nature 621, 592–601. 10.1038/s41586-023-06487-6.

11. Tortorici, M.A., Addetia, A., Seo, A.J., Brown, J., Sprouse, K., Logue, J., Clark, E., Franko, N., Chu, H., and Veesler, D. (2024). Persistent immune imprinting occurs after vaccination with the COVID-19 XBB.1.5 mRNA booster in humans. Immunity 57, 904–911 e904. 10.1016/j.immuni.2024.02.016.

12. Yisimayi, A., Song, W., Wang, J., Jian, F., Yu, Y., Chen, X., Xu, Y., Yang, S., Niu, X., Xiao, T., et al. (2024). Repeated Omicron exposures override ancestral SARS-CoV-2 immune imprinting. Nature 625, 148–156. 10.1038/s41586-023-06753-7.

13. Cao, Y., Wang, J., Jian, F., Xiao, T., Song, W., Yisimayi, A., Huang, W., Li, Q., Wang, P., An, R., et al. (2022). Omicron escapes the majority of existing SARS-CoV-2 neutralizing antibodies. Nature 602, 657–663. 10.1038/s41586-021-04385-3.

14. Lim, W.W., Mak, L., Leung, G.M., Cowling, B.J., and Peiris, M. (2021). Comparative immunogenicity of mRNA and inactivated vaccines against COVID-19. Lancet Microbe 2, e423. 10.1016/S2666-5247(21)00177-4.

15. Tas, J.M., Koo, J.-H., Lin, Y.-C., Xie, Z., Steichen, J.M., Jackson, A.M., Hauser, B.M., Wang, X., Cottrell, C.A., and Torres, J.L. (2022). Antibodies from primary humoral responses modulate the recruitment of naive B cells during secondary responses. Immunity 55, 1856–1871. e1856.

16. Schaefer-Babajew, D., Wang, Z., Muecksch, F., Cho, A., Loewe, M., Cipolla, M., Raspe, R., Johnson, B., Canis, M., and DaSilva, J. (2023). Antibody feedback regulates immune memory after SARS-CoV-2 mRNA vaccination. Nature 613, 735–742.

17. Schiepers, A., van’t Wout, M.F., Greaney, A.J., Zang, T., Muramatsu, H., Lin, P.J., Tam, Y.K., Mesin, L., Starr, T.N., and Bieniasz, P.D. (2023). Molecular fate-mapping of serum antibody responses to repeat immunization. Nature 615, 482–489.

18. Nie, J., Li, Q., Wu, J., Zhao, C., Hao, H., Liu, H., Zhang, L., Nie, L., Qin, H., Wang, M., et al. (2020). Quantification of SARS-CoV-2 neutralizing antibody by a pseudotyped virus-based assay. Nat Protoc 15, 3699–3715. 10.1038/s41596-020-0394-5.

